# Torsions-Driven Root Helical Growth, Waving And Skewing In *Arabidopsis*

**DOI:** 10.1101/2021.04.23.440761

**Authors:** Ke Zhou

## Abstract

Helical growth broadly exists in immobile plants to support their limited movement, and *Arabidopsis* seedling root exhibiting natural left-handedness helical growth is considered as a simplified model for investigating this interesting behavior. Efforts have been made for understanding the mechanism of root helical growth and consequent root waving and skewing on tilted and impenetrable surface, and several models have been established. Here, previous reports are reviewed and a straightforward torsions-driven mechanism has been emphasized, and additional experiments have been performed to fill up the gaps of this theory in our study.

This study implies that, torsions originating from handedness of both cortical microtubules and cellulose microfibrils play central role in root handed helical growth. Different from torsions directly provided by handed assembled cortical microtubules, torsions originating from right-handed assembled cellulose microfibrils are relaxed by their cross-linking with pectin within cell wall, but only exhibited when their cross-linking is interrupted due to damaged cell wall integrity. To topologically relax these torsions, supercoils of cortical microtubules and/or cellulose microfibrils exhibiting as oblique alignments are formed in root cells, which alter the orientation of root cell files and generate handed helical roots. Working together with gravitropic response, relaxation of torsions originating from helical roots drives roots to elongate with handedness, which therefore produces waved and skewed roots on tilted and impenetrable surface.

## Natural root helical growth, waving and skewing in *Arabidopsis*

Immobile plants develop various strategies to perform limited movement, among which, helical growth is one of the most efficient. Helical growth broadly exists in plants, and *Arabidopsis* root exhibiting natural but slight left-handed root helical growth provides us a simple model for understanding the mechanism of handed growth (Fig.1A) [1; 2; 3; 4]. Coupling with gel-root interaction and gravitropic response, their helical roots wave and skew with specific angles on tilted and impenetrable surface, which are surprisingly ecotype-dependent behaviors [1; 5] (Fig.1B-1D). In our study, the direction of root skewing is determined by looking downward from upside of petri-dish, which therefore opposite to our figures taken from downside of petri-dish through the medium.

**Figure.1.**
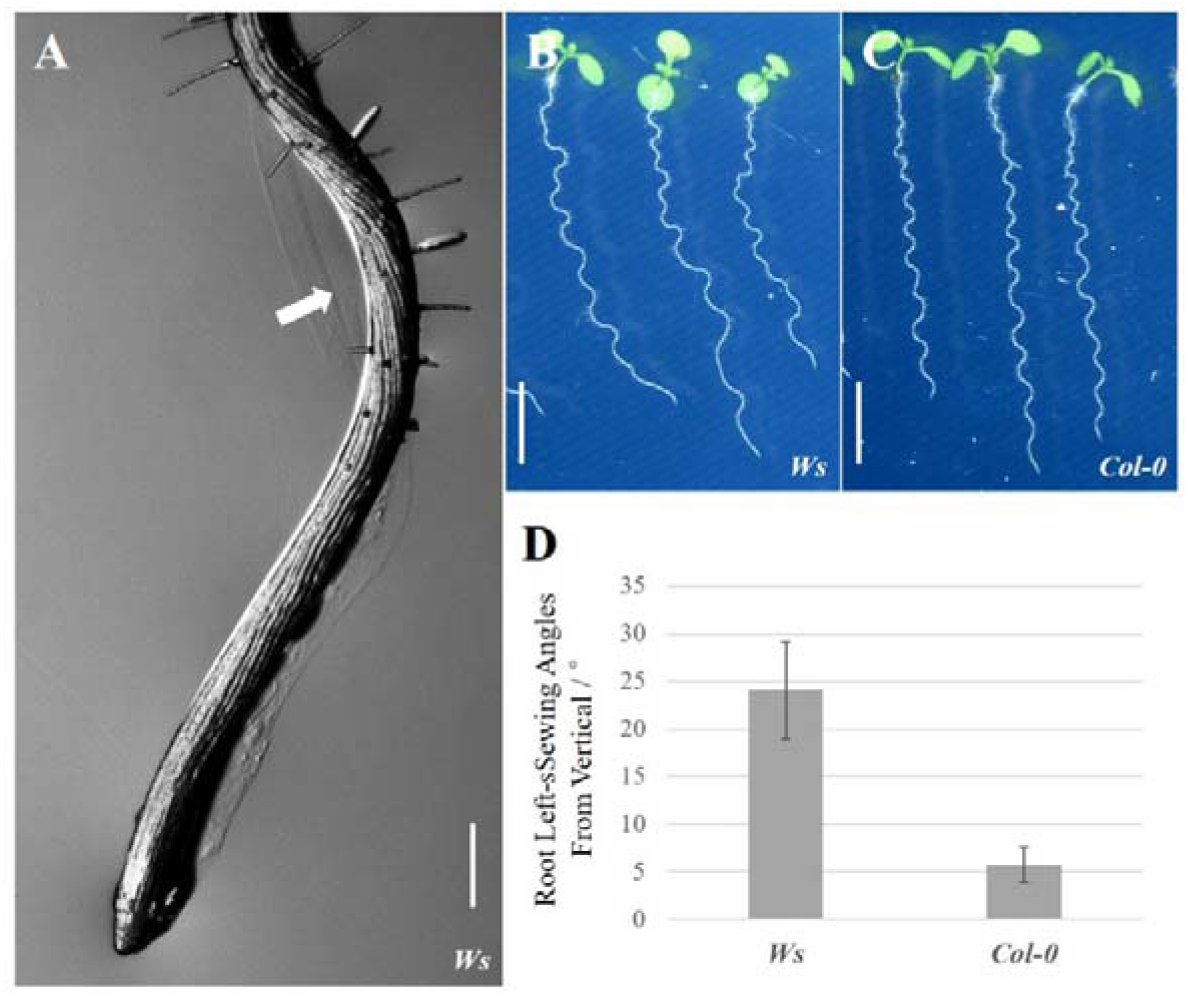
Natural root helical growth, and root waving and skewing on tilted and impenetrable surface. **(A)** Five-day-old seedling root of *Ws* ecotype *Arabidopsis*, and the helical growth was pointed by white arrow; bar, 0.2mm. Seven-day-old seedling roots from *Ws* **(B)** and col-0 **(C)** ecotype *Arabidopsis* on 30° tilted 1/2MS medium containing 1.5% agar; bar, 5mm; and their skewing angles were shown in **(D)**; n=20, and three biological repeats are performed; standard deviations are indicated as error bars..

## Cortical microtubule-determined root helical growth in *Arabidopsis*

The cortical microtubules-dependence is the most studied, and believed to be the main cause of root helical growth in *Arabidopsis* [2; 4; 6; 7; 8; 9; 10; 11; 12]. The microtubule is a hollow cylindrical structure with the diameter of 25nm generally surrounded by 13 straight protofilaments assembled by α- and β-tubulin molecules, of which typical length is several micrometers [13; 14] (Fig.2A). Both *in vitro* and *in vivo* data suggest that, microtubule from mammalian cells could be assembled by 11-16 protofilaments with a 3-4 start helix, but only microtubule with 13 protofilaments and a 3-start helix (13-3) could perfectly maintain all protofilaments parallel to the axis [15; 16; 17; 18; 19] (Fig.2C). To maintain the lattice integrity, microtubule assembled by 11-3, 12-3, 15-4, 16-4 protofilaments would cause right-handedness (Fig.2B), but 14-3 protofilaments would cause left-handedness, of which process is interestingly dynamic and reversible (Fig.2D) [15; 16; 18; 20]. Except straight microtubules with 13-3 protofilaments, left-handed microtubules with 14-3 protofilaments is predominant *in vitro* [16].

**Figure.2.**
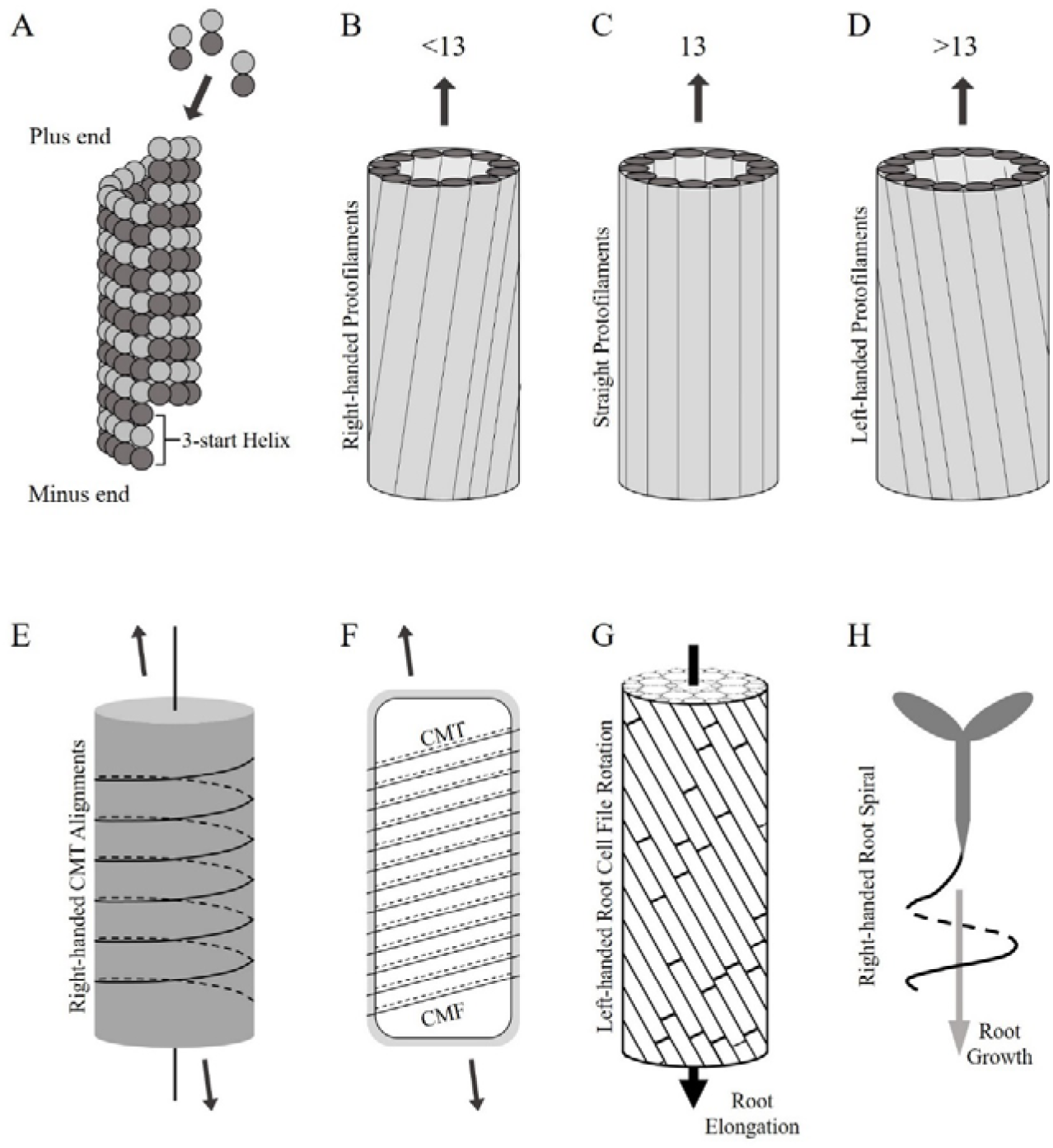
Cortical microtubule-determined root helical growth in *Arabidopsis*. **(A)**, partial typical microtubule polymerized by α- and β-tubulin molecules with 13 protofilaments and a 3-start helix; **(B)**, right-handed microtubule; **(C)**, typical microtubule and **(D)**, left-handed microtubule due to various assemble of protofilaments are shown; **(E)**, right-handed supercoil of cortical microtubule alignments; **(F)**, oblique cortical microtubules guiding cellulose microfibrils determine the direction elongation of root cell; **(G)**, left-handed root cell file rotation; and **(H)**, right-handed supercoil of root.

Shortly after mitosis, cortical microtubules are transversely aligned closely beneath to plasma membrane through an identified mechanism, which then determine cell anisotropic expansion through acting as cytoskeleton and guiding cellulose microfibrils [21; 22; 23]. If cortical microtubules with 14-3 protofilaments present in *Arabidopsis* as *in vitro*, transversely aligned left-handed cortical microtubules would exhibit right-handed supercoils to relax the torsions when stretched by the cytoplasm in rapidly elongating root cells (Fig.2E), which would be shown as obliquity of cortical microtubules (Fig.2F) [7]. The oblique cortical microtubules guiding the synthesis of cellulose microfibrils would alter the directional elongation, and form left-handed root cell file rotation (Fig.2G) which eventually cause right-handed supercoil of root (Fig.2H).

Unfortunately, direct observations on very limited number of cortical microtubules hasn’t exhibit us the presence of that with 14-3 protofilaments [8]. But considering the possibility that torsions provided by a small proportion of handed assembled cortical microtubules might be capable of re-orientating other non-handed cortical microtubules, a further quantitative study might be required for concluding or excluding the presence of left-handed cortical microtubules with 14-3 protofilaments.

## The role right-handed assembled cellulose microfibrils play in root handedness

Backbone of *Arabidopsis* root primary cell wall is composed of cellulose microfibrils cross-linking by polysaccharide molecules, such as pectin and hemicellulose [24; 25] (Fig.3B). Comparing with the uncertain handedness of cortical microtubules, cellulose microfibrils synthesized by transmembrane cellulose synthase rosettes containing six cellulose synthase are determinately right-handed assembled [2; 26] (Fig.3A). Based the same theory described above, right-handed assembled cellulose microfibrils should cause left-handed supercoil of cellulose microfibrils alignments (Fig.3D) and therefore right-handed root cell file rotation (Fig.3E).

**Figure 3.**
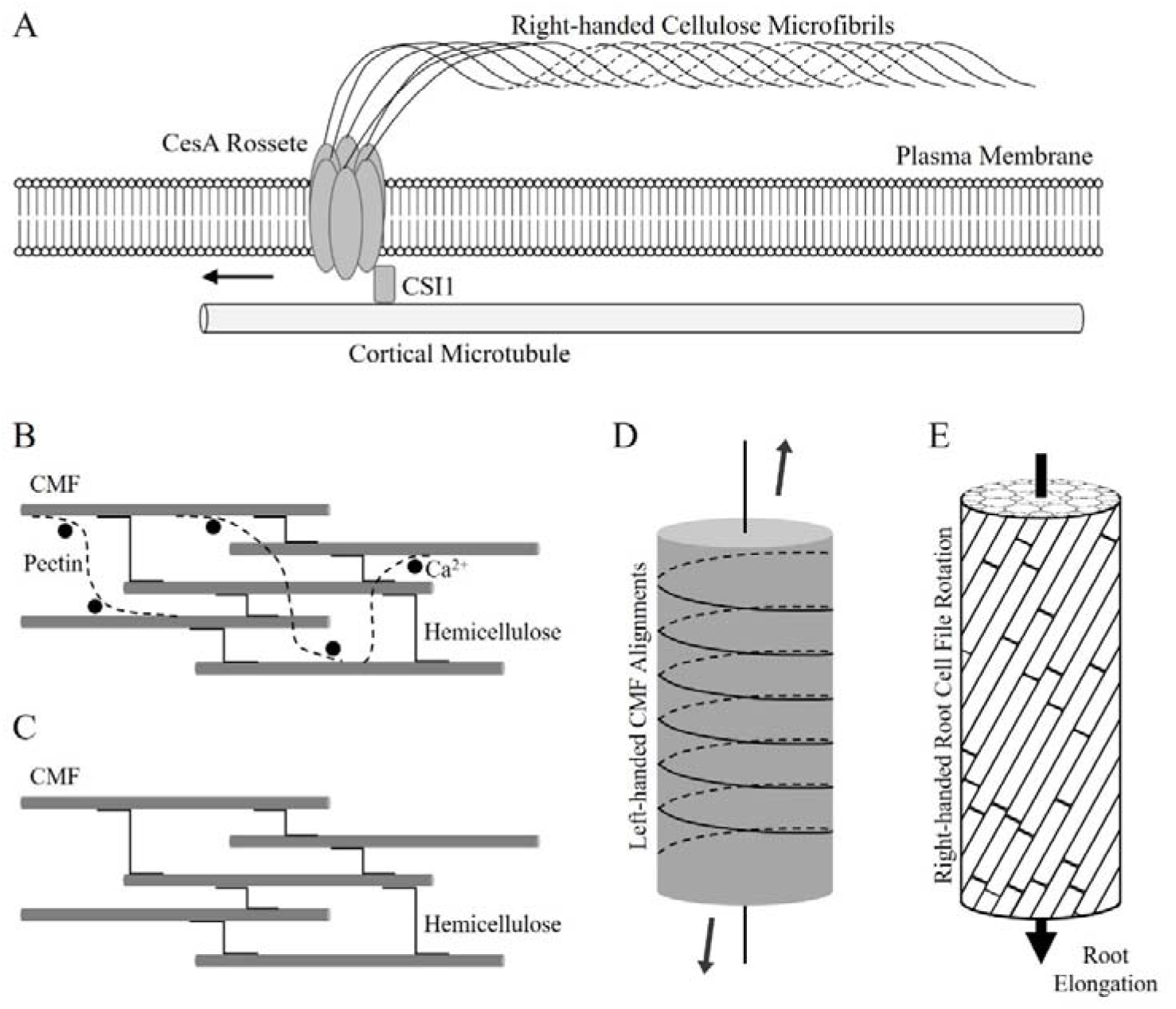
Cellulose microfibrils-determined root helical growth in *Arabidopsis*. **(A)**, the synthesis and right-handed assemble of cellulose microfibrils modified from [2]; **(B)**, cross-linked and **(C)**, decross-linked cellulose microfibrils by pectin; **(D)**, left-handed supercoil of cellulose microfibrils; and **(E)**, left-handed root cell file rotation.

But this mechanism hasn’t been clarified, whereas the cross-linking by polysaccharide molecules might be the key. Generally, cellulose microfibrils are cross-linked by flexible polysaccharide pectin [24; 25], which might be capable of relaxing the torsions originating from handed assembled cellulose microfibrils.

Actually, previous data has already exhibited us the clues: 1. Ca^2+^ is crucial for the cross-linking of cellulose microfibrils by pectin, and high concentrated NaCl displacing Ca^2+^from cell wall and therefore decross-linking cellulose microfibrils would result in right-handed root helical growth [27; 28; 29]; 2. Mutation of receptor-like kinase FER perceiving pectin decross-linked from damaged cell wall and activating Ca^2+^ export to enforce cellulose microfibrils-pectin cross-linking also results in right-handed root helical growth [30; 31]. But further experiments could clearly demonstrate this mechanism are required.

## Elongation rate-dependent root helical growth

Besides handed assembled cortical microtubules or cellulose microfibrils, another elongation rate-dependent mechanism has been revealed to be related to root helical growth in *Arabidopsis*. In *sku5* or *cob* knockout mutants, of which anisotropic cell expansion is interrupted, elongation of root cells is largely inhibited [32; 33; 34; 35; 36; 37]. But possibly due to the pressure from epidermis and cortex, defected transverse expansion of their root inner structures is inhibited, which therefore cause the different elongation rate between outer and inner structures aggravating their original helical growth [6].

## Formation of root waving and skewing on tilted and impenetrable surface

On tilted and impenetrable surface, helical root cell file rotation works together with gravitropic response to form waving and skewing roots, and several models have been established to explain their formation [1; 2; 3; 4; 38]. In this study, this process is further observed, and the traces left on medium surface by impeded point and re-localized elongation zone (Fig.4A) reveal a simplified torsion-driven model: 1, root cap fails to enter tilted medium but is impeded; 2, due to the impediment of root cap, rapid elongation drives root to curve at elongation zone, and then to left the trace on medium surface shown in Fig.4A; 3, in the process of curvature, to relax the torsions provided by the left-handed root cell file rotation at elongation zone, modified right-handed root supercoil is formed (Fig.4B); 4, as long as the gravitropic response is activated by the curvature of root tip due to right-handed root supercoil, root tip curves downward again, and repeat the processes described above to form root waving and skewing.

**Figure 4.**
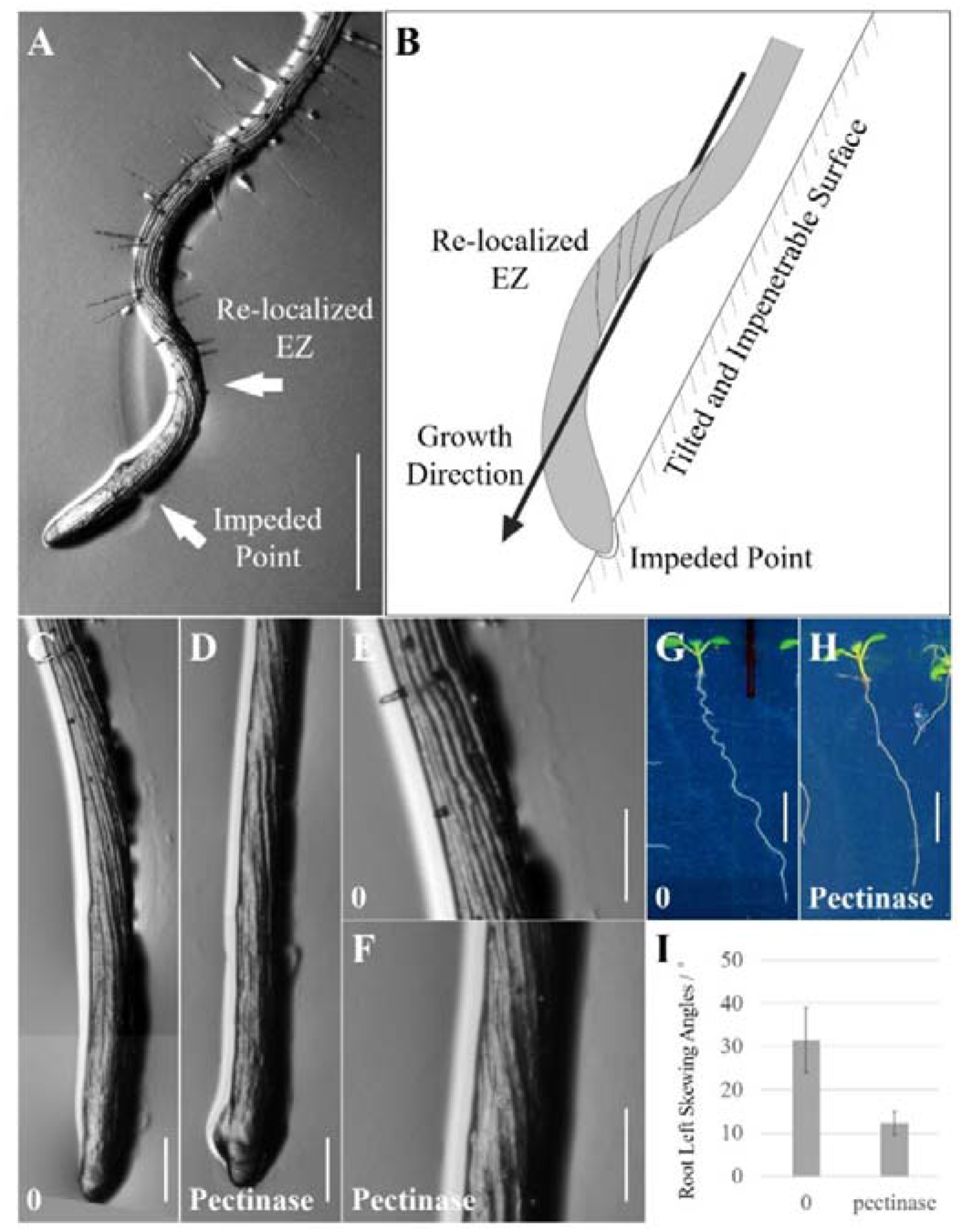
Formation of root waving and skewing, and the role cell wall play in this process. **(A)**, waved root tip from 7-day-old *Ws* ecotype *Arabidopsis* seedling by root incline assay, and traces left by impeded root cap and re-localized elongation zone are pointe by white arrows; bar,1mm; **(B)**, formation of right-handed root supercoil originating from impedition of root cap and rapid elongation of root at elongation zone; 7-day-old *Ws* ecotype *Arabidopsis* root tips by root incline assay without **(C** and **F)** or with pectinase **(D** and **G)** are shown; bar,0.5mm; their root waving and skewing are shown in **(E)** and **(F)**; bar,5mm; and the skewing angles are indicated in **(G)**; n=20, and error bars are indicated.

In this process, cell wall mediates the relaxation of torsions, and application of pectinase solution from *Aspergillus niger* (P4716-Sigma-aldrich) reducing cell wall mechanic strength through digesting cell wall components strongly suppresses root waving and skewing (Fig.4G-4I), although their root helical growth is not significantly affected (Fig.4C-4F).

## The role gravitropic response plays in root waving and skewing

As described above, gravitropic response works to re-orientate the curved root tips due to relaxation of torsion from left-handed root cell file rotation, and to form root waving. Therefore, application of NPA blocking auxin polar transport and suppressing tropic response would significantly suppressing the root waving, but form root supercoils (Fig.5A-5C).

**Figure 5.**
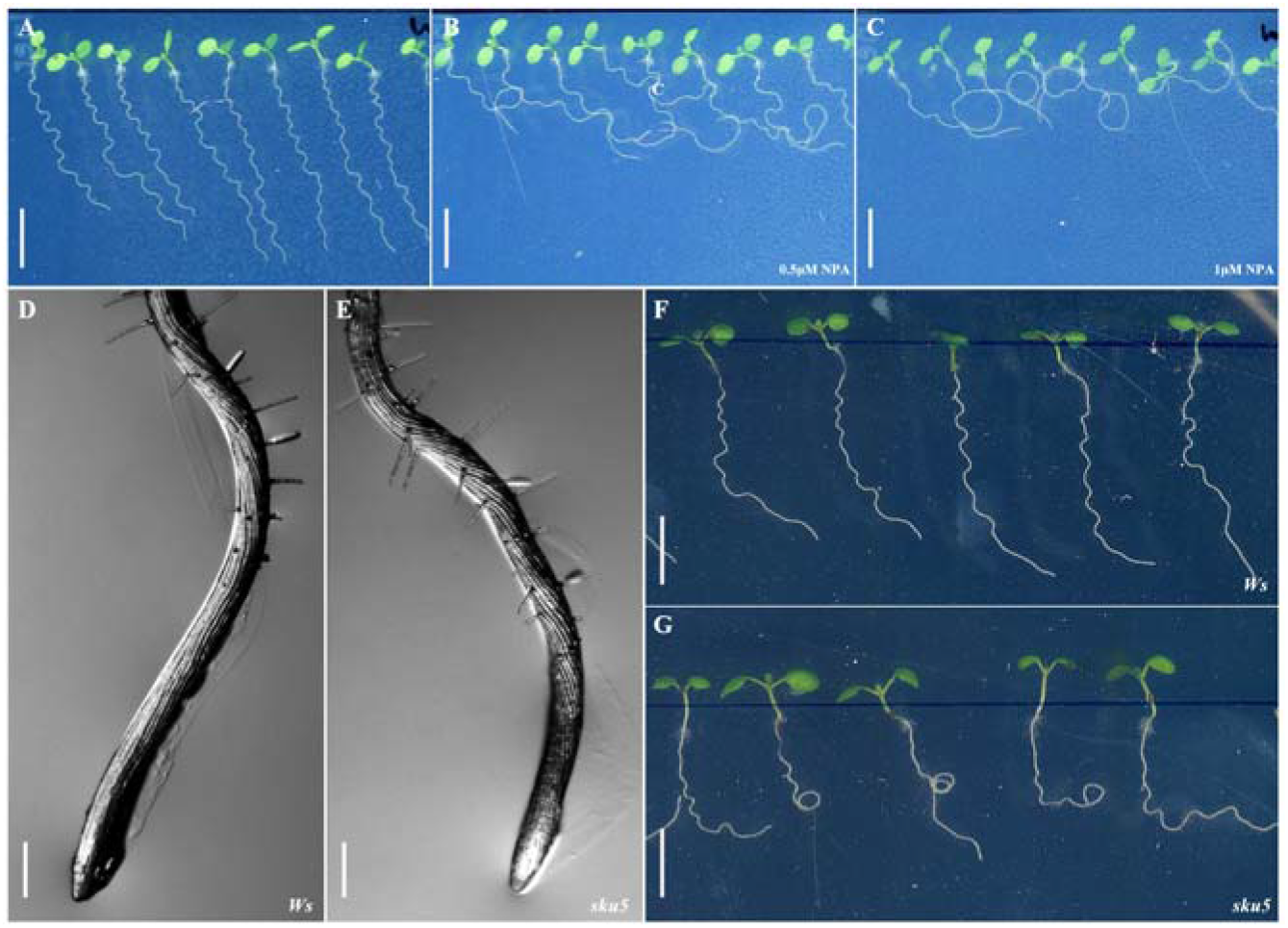
Gravitropic response in formation of root waving and skewing. Incline growth assay is utilized on *Ws* ecotype *Arabidopsis* without **(A)**, or with 0.5μM **(B)** and 1μM **(C)** NPA blocking auxin polar transport; bar,0.5cm. Root tips of 5-day-old seedlings from **(D)** *Ws* and **(E)** *sku5-2* mutant allele, bar=0.5mm; and their root waving and skewing by incline growth assay are shown in **(F)** and **(G)**; bar,0.5cm.

On the contrary, *sku5* mutant exhibiting more severe left-handedness of root cell file rotation without altering gravitropic response [32] (Fig.5D-5E) would overcome the influence from gravitropic response, and also form root supercoils (Fig.5F-5G).

It implies that both tropic response and helical growth participate in formation of root waving and skewing, which make it unreliable to determine the root helical growth simply through observing root waving and skewing, especially in those mutants affecting auxin polar distribution [39; 40; 41].

## DISCUSSION

Our study emphasizes a useful and simplified mechanism to understand the helical behaviors of *Arabidopsis* roots which might also work on other organs and species. This model could be supported by the root helical behaviors found in various *Arabidopsis* mutants which have been well reviewed recently in [2] and re-classified according to the origins mentioned in our study: Class 1, mutations on tubulin molecules or MAPs altering handedness of cortical microtubules, such as *lefty1* and *lefty2* alleles modifying α-tubulin6 and α-tubulin4 respectively [9], and *loss-of-function* of a microtubule plus-end tracking protein *SPR1* [11; 42], which cause respective helical behaviors; Class 2, *loss-of-functions* of cell wall integrity-related components, such as *FER* perceiving and recovering damaged cell wall integrity, which result in right-handed helical behaviors [30; 31]; Class 3, *loss-of-functions* of genes regulating cell anisotropic expansion, such as *COB* [35] and *SKU5* [32], which aggravate original helical behaviors; and Class 4, *loss-of-functions* of genes regulating auxin polar distribution, such as *AUX1* [39; 40; 43], *TWD1* [44] and *RCN1* [41; 45], which do not alter root helical growth, but alter other behaviors such as root waving and skewing. To understand the mechanisms of altered root helical behaviors in other mutants, further investigations are required.

## PERSPECTIVES

To complete the model of torsions-driven helical behaviors in *Arabidopsis* root, urgent tasks are required: 1, quantitative data of the dynamic cortical microtubules assembled by different numbers of protofilaments; 2, convincing evidence of helical behaviors originating from right-handed assembled cellulose microfibrils.

## METHODS

### Plant materials

*T-DNA* insertion line *sku5-2* (FLAG_300H05) is in *Ws* ecotype *Arabidopsis* background.

### Incline growth assay

Seeds of *Ws* and *sku5-2* mutant are sowed on 30° tilted 1/2MS medium containing 1.5% Agar, and then grow in green chamber under 16/8 hours light at 22°C for 7 days. In our study, the direction of root skewing is determined by looking downward from upside of petri-dish, which therefore opposite to our figures taken from downside of petri-dish through the medium.

### Pectinase treatment

1mM pectinase solution from *Aspergillus niger* (P4716-Sigma-aldrich), which could reduce cell wall mechanic strength through digesting cell wall components, is added in 1L cooled 1/2MS medium, and then poured into petri-dishes.

## References

[1] R. Roy, and D.C. Bassham, Root growth movements: waving and skewing. Plant Sci 221-222 (2014) 42–7.

[2] H. Buschmann, and A. Borchers, Handedness in plant cell expansion: a mutant perspective on helical growth. New Phytol 225 (2020) 53–69.

[3] F. Migliaccio, and S. Piconese, Spiralizations and tropisms in Arabidopsis roots. Trends Plant Sci 6 (2001) 561–5.

[4] D.R. Smyth, Helical growth in plant organs: mechanisms and significance. Development 143 (2016) 3272–82.

[5] F. Migliaccio, A. Fortunati, and P. Tassone, Arabidopsis root growth movements and their symmetry: progress and problems arising from recent work. Plant Signal Behav 4 (2009) 183–90.

[6] T. Ishida, S. Thitamadee, and T. Hashimoto, Twisted growth and organization of cortical microtubules. J Plant Res 120 (2007) 61–70.

[7] L.M. Vaughn, K.L. Baldwin, G.X. Jia, J.C. Verdonk, A.K. Strohm, and P.H. Masson, The Cytoskeleton and Root Growth Behavior. Advan Pl Bio 2 (2011) 307–326.

[8] T. Ishida, Y. Kaneko, M. Iwano, and T. Hashimoto, Helical microtubule arrays in a collection of twisting tubulin mutants of Arabidopsis thaliana. Proc Natl Acad Sci U S A 104 (2007) 8544–9.

[9] S. Thitamadee, K. Tuchihara, and T. Hashimoto, Microtubule basis for left-handed helical growth in Arabidopsis. Nature 417 (2002) 193–196.

[10] J.C. Sedbrook, and D. Kaloriti, Microtubules, MAPs and plant directional cell expansion. Trends Plant Sci 13 (2008) 303–10.

[11] I. Furutani, Y. Watanabe, R. Prieto, M. Masukawa, K. Suzuki, K. Naoi, S. Thitamadee, T. Shikanai, and T. Hashimoto, The SPIRAL genes are required for directional control of cell elongation in Aarabidopsis thaliana. Development 127 (2000) 4443–53.

[12] J.C. Sedbrook, D.W. Ehrhardt, S.E. Fisher, W.R. Scheible, and C.R. Somerville, The Arabidopsis sku6/spiral1 gene encodes a plus end-localized microtubule-interacting protein involved in directional cell expansion. Plant Cell 16 (2004) 1506–20.

[13] M. Knossow, V. Campanacci, L.A. Khodja, and B. Gigant, The Mechanism of Tubulin Assembly into Microtubules: Insights from Structural Studies. iScience 23 (2020) 101511.

[14] G. Duclos, R. Adkins, D. Banerjee, M.S.E. Peterson, M. Varghese, I. Kolvin, A. Baskaran, R.A. Pelcovits, T.R. Powers, A. Baskaran, F. Toschi, M.F. Hagan, SJ. Streichan, V. Vitelli, D.A. Beller, and Z. Dogic, Topological structure and dynamics of three-dimensional active nematics. Science 367 (2020) 1120–1124.

[15] D. Chretien, and R.H. Wade, New data on the microtubule surface lattice. Biol Cell 71 (1991) 161–74.

[16] G.B. Pierson, P.R. Burton, and R.H. Himes, Alterations in number of protofilaments in microtubules assembled in vitro. J Cell Biol 76 (1978) 223–8.

[17] N. Yagi, S. Matsunaga, and T. Hashimoto, Insights into cortical microtubule nucleation and dynamics inArabidopsisleaf cells. Journal of Cell Science 131 (2018) jcs203778.

[18] H. Sui, and K.H. Downing, Structural basis of interprotofilament interaction and lateral deformation of microtubules. Structure 18 (2010) 1022–31.

[19] R. Dallai, P. Lupetti, and C. Mencarelli, Unusual axonemes of hexapod spermatozoa. Int Rev Cytol 254 (2006) 45–99.

[20] M. Nakamura, and T. Hashimoto, Mechanistic Insights into Plant Chiral Growth. Symmetry 12 (2020) 2056.

[21] A.R. Paredez, C.R. Somerville, and D.W. Ehrhardt, Visualization of cellulose synthase demonstrates functional association with microtubules. Science 312 (2006) 1491–5.

[22] C.T. Anderson, A. Carroll, L. Akhmetova, and C. Somerville, Real-time imaging of cellulose reorientation during cell wall expansion in Arabidopsis roots. Plant Physiol 152 (2010) 787–96.

[23] D.J. Cosgrove, Nanoscale structure, mechanics and growth of epidermal cell walls. Curr Opin Plant Biol 46 (2018) 77–86.

[24] E.R. Lampugnani, G.A. Khan, M. Somssich, and S. Persson, Building a plant cell wall at a glance. J Cell Sci 131 (2018).

[25] A. Peaucelle, S. Braybrook, and H. Hofte, Cell wall mechanics and growth control in plants: the role of pectins revisited. Front Plant Sci 3 (2012) 121.

[26] S. Rongpipi, D. Ye, E.D. Gomez, and E.W. Gomez, Progress and Opportunities in the Characterization of Cellulose - An Important Regulator of Cell Wall Growth and Mechanics. Front Plant Sci 9 (2018) 1894.

[27] T. Shoji, K. Suzuki, T. Abe, Y. Kaneko, H. Shi, J.K. Zhu, A. Rus, P.M. Hasegawa, and T. Hashimoto, Salt stress affects cortical microtubule organization and helical growth in Arabidopsis. Plant Cell Physiol 47 (2006) 1158–68.

[28] C.S. Byrt, R. Munns, R.A. Burton, M. Gilliham, and S. Wege, Root cell wall solutions for crop plants in saline soils. Plant Sci 269 (2018) 47–55.

[29] H.W. Koyro, Ultrastructural and physiological changes in root cells of Sorghum plants (Sorghum bicolor x S. sudanensis cv. Sweet Sioux) induced by NaCl. Journal of Experimental Botany 48 (1997) 693–706.

[30] H.W. Shih, N.D. Miller, C. Dai, E.P. Spalding, and G.B. Monshausen, The receptor-like kinase FERONIA is required for mechanical signal transduction in Arabidopsis seedlings. Curr Biol 24 (2014) 1887–92.

[31] W. Feng, D. Kita, A. Peaucelle, H.N. Cartwright, V. Doan, Q. Duan, M.C. Liu, J. Maman, L. Steinhorst, I. Schmitz-Thom, R. Yvon, J. Kudla, H.M. Wu, A.Y. Cheung, and J.R. Dinneny, The FERONIA Receptor Kinase Maintains Cell-Wall Integrity during Salt Stress through Ca(2+) Signaling. Curr Biol 28 (2018) 666–675 e5.

[32] J.C. Sedbrook, K.L. Carroll, K.F. Hung, P.H. Masson, and C.R. Somerville, The Arabidopsis SKU5 gene encodes an extracellular glycosyl phosphatidylinositol-anchored glycoprotein involved in directional root growth. Plant Cell 14 (2002) 1635–48.

[33] K. Zhou, GPI-anchored SKU5/SKS are maternally required for integument development in Arabidopsis. biorxiv (2019).

[34] G. Schindelman, A. Morikami, J. Jung, T.I. Baskin, N.C. Carpita, P. Derbyshire, M.C. McCann, and P.N. Benfey, COBRA encodes a putative GPI-anchored protein, which is polarly localized and necessary for oriented cell expansion in Arabidopsis. Genes Dev 15 (2001) 1115–27.

[35] F. Roudier, A.G. Fernandez, M. Fujita, R. Himmelspach, G.H. Borner, G. Schindelman, S. Song, T.I. Baskin, P. Dupree, G.O. Wasteneys, and P.N. Benfey, COBRA, an Arabidopsis extracellular glycosyl-phosphatidyl inositol-anchored protein, specifically controls highly anisotropic expansion through its involvement in cellulose microfibril orientation. Plant Cell 17 (2005) 1749–63.

[36] D. Ben-Tov, Y. Abraham, S. Stav, K. Thompson, A. Loraine, R. Elbaum, A. de Souza, M. Pauly, JJ. Kieber, and S. Harpaz-Saad, COBRA-LIKE2, a member of the glycosylphosphatidylinositol-anchored COBRA-LIKE family, plays a role in cellulose deposition in arabidopsis seed coat mucilage secretory cells. Plant Physiol 167 (2015) 711–24.

[37] E. Niu, X. Shang, C. Cheng, J. Bao, Y. Zeng, C. Cai, X. Du, and W. Guo, Comprehensive Analysis of the COBRA-Like (COBL) Gene Family in Gossypium Identifies Two COBLs Potentially Associated with Fiber Quality. PLoS One 10 (2015) e0145725.

[38] M.V. Thompson, and N.M. Holbrook, Root-gel interactions and the root waving behavior of Arabidopsis. Plant Physiol 135 (2004) 1822–37.

[39] K. Okada, and Y. Shimura, Reversible root tip rotation in Arabidopsis seedlings induced by obstacle-touching stimulus. Science 250 (1990) 274–6.

[40] J.I. Mirza, The Effects of Light and Gravity on the Horizontal Curvature of Roots of Gravitropic and Agravitropic Arabidopsis thaliana L. Plant Physiol 83 (1987) 118–20.

[41] J. Deruere, K. Jackson, C. Garbers, D. Soll, and A. Delong, The RCN1-encoded A subunit of protein phosphatase 2A increases phosphatase activity in vivo. Plant J 20 (1999) 389–99.

[42] C. Galva, V. Kirik, J.J. Lindeboom, D. Kaloriti, D.M. Rancour, P.J. Hussey, S.Y. Bednarek, D.W. Ehrhardt, and J.C. Sedbrook, The microtubule plus-end tracking proteins SPR1 and EB1b interact to maintain polar cell elongation and directional organ growth in Arabidopsis. Plant Cell 26 (2014) 4409–25.

[43] R. Swarup, and R. Bhosale, Developmental Roles of AUX1/LAX Auxin Influx Carriers in Plants. Front Plant Sci 10 (2019) 1306.

[44] M. Wildwater, A. Campilho, J.M. Perez-Perez, R. Heidstra, I. Blilou, H. Korthout, J. Chatterjee, L. Mariconti, W. Gruissem, and B. Scheres, The RETINOBLASTOMA-RELATED gene regulates stem cell maintenance in Arabidopsis roots. Cell 123 (2005) 1337–49.

[45] M. Karampelias, P. Neyt, S. De Groeve, S. Aesaert, G. Coussens, J. Rolcik, L. Bruno, N. De Winne, A. Van Minnebruggen, M. Van Montagu, M.R. Ponce, J.L. Micol, J. Friml, G. De Jaeger, and M. Van Lijsebettens, ROTUNDA3 function in plant development by phosphatase 2A-mediated regulation of auxin transporter recycling. Proc Natl Acad Sci U S A 113 (2016) 2768–73.

